# Hierarchical recurrent temporal prediction as a model of the mammalian dorsal visual pathway

**DOI:** 10.1101/2025.05.15.654214

**Authors:** Sebastian Klavinskis-Whiting, Andrew J. King, Nicol S. Harper

## Abstract

A major goal of neuroscience is to identify whether there are generalized principles that can explain the diverse structures and functions of the brain. The principle of temporal prediction provides one approach, arguing that the sensory brain is optimized to represent stimulus features that efficiently predict the immediate future input. Previous work has demonstrated that feedforward hierarchical temporal prediction models can capture the tuning properties of neurons along the visual pathway, and that recurrent temporal prediction models can explain local functional connectivity within primary visual cortex. However, the visual system is also characterized by extensive inter-areal feedback recurrency, which existing models lack. We aimed to better account for the dynamic features of neurons in the visual cortex by incorporating both local recurrency and inter-areal feedback connectivity into a hierarchical temporal prediction model. The resulting model captured tuning for pattern motion, surround suppression and elements of inter-areal functional connectivity in visual cortex. Moreover, compared with several alternative normative models, the hierarchical recurrent temporal prediction model provided the closest fit to these tuning properties and was best able to explain the emergence of neuronal response properties across the visual cortex. Accordingly, temporal prediction can account for information processing throughout the visual pathway.

**Author summary:** Task-driven artificial neural networks have provided valuable insights into the computational operations performed by neurons in the brain. In particular, models trained on various normative principles have been shown to capture some of the feature selectivity of neurons in the visual system. However, these models generally ignore the substantial role of inter-areal feedback connectivity in neural function. This study presents an alternative approach to understanding the neural computations underlying the functional organization of the visual system, by providing compelling evidence that networks trained for temporal prediction can explain the emergence of higher-level response properties across the visual cortex and relating these features to both local and inter-areal feedback connectivity.

## Introduction

Classical theories of vision posit a series of hierarchically organized processing stages that gradually extract higher-level visual features [1,2]. In this way, the brain is thought to decompose the patterns of light that impinge on the retina into meaningful representations to be further processed in downstream areas, and ultimately, to guide behavior. A key question, then, is how to understand these computations and whether an underlying principle of organization can explain how the resulting representations change across the visual pathway [3,4].

One promising principle is that of temporal prediction, which argues that the sensory brain is optimized to represent stimulus features that are predictive of the immediate future [4–7]. This is likely to be useful for extracting underlying variables in the input, eliminating unnecessary information based on the behavioral relevance of stimuli [8], and creating a representation that is useful for guiding future action given sensory and motor delays [4–6]. Temporal prediction can be applied in a hierarchical manner to capture spatiotemporal receptive field properties across multiple stages of the visual pathway [4]. Importantly, as an unsupervised principle, temporal prediction does not rely on hand-labeled examples, as in many deep learning models of the visual system [9,10]. Together, these features make temporal prediction a promising candidate for modeling neural processing in sensory systems.

In addition to demonstrating that the tuning properties of neurons along the visual pathway can be reproduced by applying temporal prediction in a hierarchical manner [4,5], we have recently shown that the functional specificity of connections within the primary visual cortex (V1) can be captured by applying temporal prediction to a locally-recurrent network [6]. However, these models have so far neglected the substantial role of long-range feedback connectivity from higher-order visual areas, which has been implicated in a range of visual processing functions [11]. Feedback connectivity in the visual system appears to impart both modulatory and driver roles on downstream neural activity [11,12]. Among other roles, feedback from higher visual areas is believed to mediate extra-classical receptive field effects, such as surround suppression [13–16], maintain and gate working memory representations [17–19], and, within the predictive coding framework, convey predictions that may drive learning via bottom-up prediction errors [20,21]. Accordingly, incorporating these elements into normative models, including temporal prediction, is likely to be an important step in explaining the structural and functional properties of the visual pathway.

Here, we demonstrate that a hierarchical recurrent temporal prediction model trained on movies of dynamic natural scenes better captures the tuning properties of visual neurons along the mammalian visual pathway than implementing hierarchy or local recurrency alone. The architectural addition of inter-areal recurrency and the network’s optimization for temporal prediction jointly improved the model’s fit to the tuning properties of visual cortical neurons measured in different studies. Moreover, feedback connectivity significantly improved the network’s memory capacity and frame prediction performance, and recapitulated the dependence on feedback of surround suppression in the responses

of V1 neurons [13–16]. Finally, compared with alternative normative models, the hierarchical recurrent temporal prediction model was more closely aligned with the stimulus representations that have been described in different areas of the visual cortex. Together, these results provide evidence that inter-areal feedback across the visual hierarchy may also be optimized for temporal prediction.

## Results

### The hierarchical recurrent model improves temporal prediction performance

The hierarchical recurrent model consisted of a recurrent neural network instantiating a hierarchical model of temporal prediction, where each group of units was trained to predict its lower-order inputs. Thus, the first group (G_1_) was trained for next-frame prediction of video clips of dynamic natural scenes, while the second and third groups (G_2_ and G_3_) were trained to predict the future activity of groups G_1_ and G_2_, respectively (Fig 1A). Within the recurrent layer of the network, internal connectivity ensured that each group received ‘feedforward’ input from the previous group, ‘feedback’ input from the subsequent group and local recurrent input from itself. We conceptualized each group as modeling increasingly deep regions of the dorsal visual pathway, with groups G_1_, G_2_, and G_3_ in particular corresponding to primary visual cortex (V1), secondary visual cortex (V2) and middle temporal area (MT), respectively (Fig 1B). However, the model structure and training data are not *per se* tailored to any specific mammalian species. Hence, while we primarily compare our model to macaque data, where such data are not available, we compare the model to mouse data.

**Fig 1.**
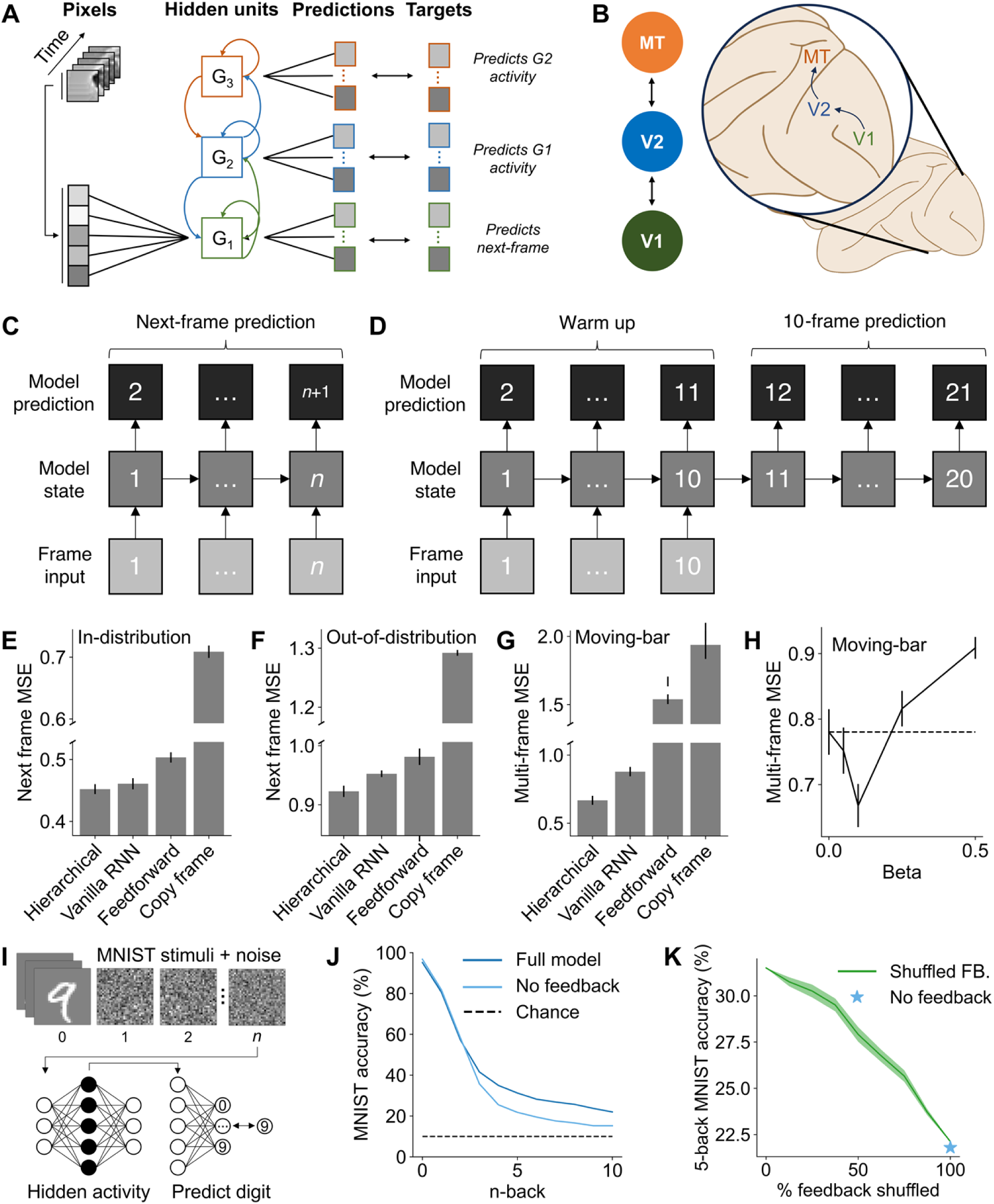
The hierarchical recurrent temporal prediction model shows improved frame prediction performance. (A) Schematic of the temporal prediction model. (B) A subset of the dorsal visual pathway, with regions V1, V2 and MT highlighted, putatively corresponding to groups G_1_, G_2_ and G_3_. (C) The next frame prediction procedure where the preceding frame and hidden state are inputted into the model at each timestep. (D) The multi-frame prediction procedure, where the model was run autoregressively such that the model predicted multiple frames, relying on its own predictions and hidden state alone. (E-G) Prediction performance across comparison models for (E) in-distribution, (F) out-of-distribution and (G) multi-frame prediction of a moving bar. In all cases, the hierarchical model performed best, exceeding the performance of the comparison models. (H) Multi-frame prediction performance for a moving bar across differing beta values. The lowest mean squared error was found for a beta value of 0.1, implying that a hierarchical instantiation of the network can improve prediction performance. (I) Schematic of the n-back procedure. MNIST digits are used as input to the model followed by *n* frames of Gaussian noise. The hidden state is then linearly mapped to predict the digit identity to calculate an accuracy score, repeating the same procedure for all *n* 0-10. (J) MNIST accuracy declines more quickly when feedback is ablated, suggesting that model feedback increases the memory capacity of the network. (K) MNIST accuracy declines as the percentage of shuffled feedback weights increases, implying that the increase in memory capacity is not purely a feature of the model’s architecture, but depends on the trained weights of the feedback connectivity.

We first analyzed the temporal prediction performance of the hierarchical recurrent model and compared this to several baseline models. Specifically, we compared the hierarchical recurrent temporal prediction model to a single-layer recurrent temporal prediction model without the addition of hierarchical groups, a purely feedforward model that omitted both local and feedback recurrency, and a baseline comparison where we compared the predictions to simply copying the input frame. The copy frame comparison ensured that each model was truly acting as a generative model to predict the next frame, rather than merely reproducing its inputs as a trivial solution. For next frame prediction (Fig 1C), we calculated the mean squared error between the true and predicted next frame of the stimulus. Conversely, for 10 frame prediction (Fig 1D), we measured the mean squared error across 10 frames generated in an autoregressive manner. This involved repeatedly predicting the next frame using the model’s own preceding predictions, rather than the true frame inputs. For next frame prediction, we assessed the model’s performance both in-distribution (the held-out test set from the stimuli used for training; Fig 1E) and out-of-distribution (a novel dataset to that used during training; Fig 1F) to assess the generality of these findings. In the multi-frame case, the input consisted of oriented moving bar stimuli where there was an unambiguous ‘true future’ given the preceding input (Fig 1G, H).

Across all conditions, we found that the hierarchical model performed best, with the overall lowest mean squared error for temporal prediction (paired t-test, all *p*<0.0001) (Fig 1E-G). Thus, the addition of hierarchy in the form of additional groups improved prediction performance compared with both a single-layer recurrent model and a feedforward model trained for temporal prediction. Finally, all three models exceeded the baseline performance for copying the preceding frame (paired t-test, all *p*<0.0001).

We also varied the beta parameter, which controls the influence of higher groups on the hierarchical recurrent model’s loss function during training, to assess its impact on temporal prediction performance (Fig 1H). A higher beta value indicates a greater weighting applied to higher groups, with β=0.5 indicating an equal weighting. Although higher beta values impaired next-frame prediction performance across the in- and out-of-distribution cases (Fig S1), we found that a small beta improved multi-frame prediction on the moving bar stimuli set (β=0 vs β=0.1: paired t-test, *t*(71)=3.20, *p*=0.0021). Thus, the addition of hierarchy to the loss function could improve the model’s capacity to generalize for multi-frame prediction across novel stimuli.

Given the importance of hierarchy in the model for temporal prediction, we next asked whether this might in part be supported by increasing the memory capacity of the model, mediated by the network’s feedback connectivity. In the visual cortex, feedback from higher visual areas to V1 is selectively recruited during working memory tasks [18] while disruption of corticocortical feedback to higher visual areas has been shown to impair performance in working memory-dependent behavior [19]. Thus, we reasoned that feedback in the hierarchical recurrent temporal prediction model might similarly support the network’s memory capacity.

To test this hypothesis, we trained a linear decoder to predict the identity of hand-written MNIST digits from the network’s hidden state. To probe how well the model’s hidden state could maintain the digit representations, the decoder was trained based on the hidden state after *n* frames of Gaussian noise input (Fig 1I). For each n-back decoder, we then calculated the decoder’s accuracy on the MNIST test set for the model with and without feedback. Ablating feedback significantly reduced the decoder accuracy from 3-back onwards (z-test, *z=*6.54*, p*<0.0001), implying that feedback improved the model’s capacity to maintain representations over longer timescales (Fig 1J). To test whether this improvement was related to the model’s training or whether any form of feedback could account for the difference, we assessed the 5-back accuracy when shuffling an increasing proportion of the model’s feedback weights (Fig 1K). As the proportion of shuffled weights increased, the accuracy decreased until it was non-significantly different from the model where feedback was ablated entirely (z-test, *z*=0.494*, p*=0.621). Thus, randomized feedback was ineffective, and feedback weights required the structure imposed by training to maintain network representations.

### Hierarchical recurrent temporal prediction captures tuning properties across the visual hierarchy

We next investigated how model unit response properties varied across the model’s groups relative to different stages of the visual system. To make these comparisons, we estimated the model units’ receptive fields using the response-weighted average of each unit to Gaussian random noise. Units in the first group generally had a clear Gabor-like structure with alternating excitatory and inhibitory regions akin to the receptive fields of V1 simple cells (Fig 2Ai). In contrast, units in higher-order groups generally displayed little spatial structure (Fig 2Aii, iii), consistent with a more complex-cell-like response that is not well estimated by the response-weighted averaging procedure [22].

**Fig 2.**
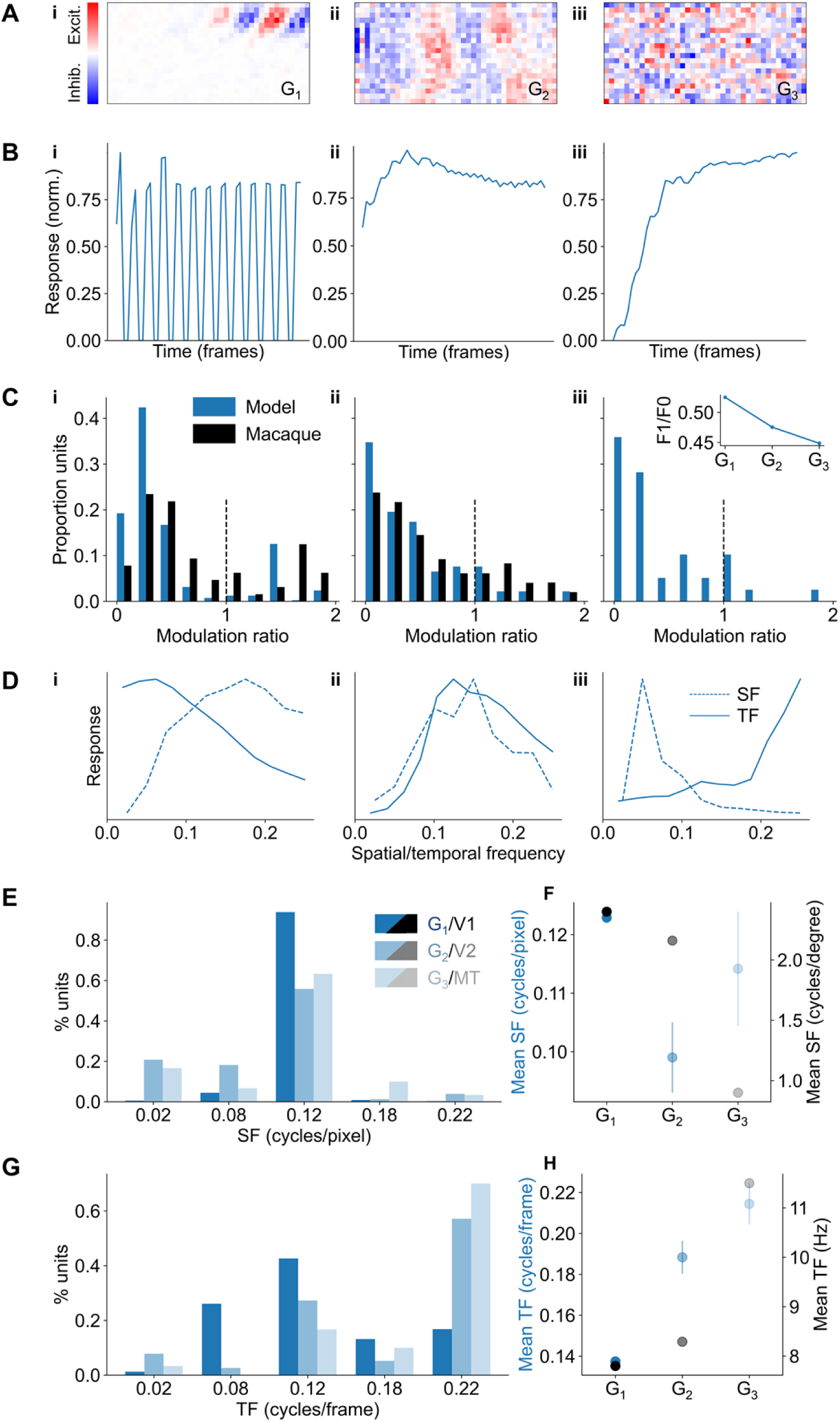
Hierarchical recurrent temporal prediction captures tuning properties across the visual hierarchy. (A) Exemplar model unit receptive fields from group G_1_ (i), G_2_ (ii) and G_3_ (iii). (B) Normalized response to the preferred grating stimulus for the same model units as in (A). (C) Distribution of modulation ratio values across model groups G_1_ (i), G_2_ (ii) and G_3_ (iii) (blue bars). Comparison data for G_1_ from macaque V1 and for G_2_ from macaque V2 [25] (black bars) are also shown. Inset shows the average modulation value for each model group. (D) Exemplar spatial and temporal frequency tuning curves for three units from G_1_ (i), G_2_ (ii) and G_3_ (iii). (E) Distribution of preferred spatial frequencies for model units across each group. (F) Mean preferred spatial frequency for each group compared with macaque V1 [26], V2 [27], and MT [28]. (G) Distribution of preferred temporal frequencies for model units across each group. (H) Mean preferred temporal frequency for each group compared with macaque V1 [28], V2 [27], and MT [28].

To further characterize the receptive field properties of model units, we recorded their responses to full-field sinusoidal gratings of varying temporal frequency, spatial frequency, and orientation. The preferred stimulus was defined as the drifting grating that maximally stimulated each unit, which we used to classify model units as simple-cell- or complex-cell-like in their responses [22,23]. In visual cortex, the activity of simple cells is heavily modulated by a drifting grating, whereas complex-cell responses are phase invariant and only minimally modulated by the moving stimulus. We found model units that exhibited both simple-cell- (Fig 2Bi) and complex-cell-like (Fig 2Bii, 2Biii) responses.

We quantified these responses using the modulation ratio – the ratio of their amplitudes to that of the average response – with a larger modulation ratio indicating a more simple-cell-like response (Fig 2C). G_1_ units (mean modulation ratio=0.53) had a bimodal-like distribution similar to that of macaque V1 [24] (mean=0.79), indicating two subpopulations of units (Fig 2Ci). Higher groups (G_2_ mean=0.48, G_3_ mean=0.45) had a more unimodal-like distribution, which tended to monotonically decline with increasing modulation ratio, similarly to the true distribution found in V2 [25] (mean=0.63) (Fig 2Cii, 2Ciii). This was reflected in the average modulation ratio across groups (Fig 2Ciii inset), which declined for higher model groups, indicating a larger proportion of complex-cell-like responses. Finally, we calculated the joint distributions of the modulation ratio with the model unit’s orientation selectivity (Fig S2A), as well as how well each unit was modelled as a Gabor (Fig S2B). As expected, and in tandem with the biology, simple-cell-like model units were more orientation selective (mean orientation selectivity index, OSI=0.84) and had a higher *r^2^* for their Gabor fits (mean=0.60) than complex-cell-like units (mean OSI=0.77; t-test, *t*(181)=-3.84, *p*=0.0002; mean Gabor fit=0.54 t-test, *t*(251)=-4.23, *p*<0.0001).

The distribution of spatial and temporal frequency tuning properties also varied systematically across the model. For each unit, we measured the tuning curve for that unit’s response as a function of temporal and spatial frequency, with the preferred frequency taken as the tuning curve’s peak. Units were generally well tuned, with tuning curves spanning a wide range of temporal and spatial frequencies (Fig 2D, 2E). The second group G_2_ had a lower average preferred spatial frequency than the first group G_1_ (t-test, *t*(79.4)=3.91, *p*=.0002; Fig 2F). The same trend has been found in macaque visual cortex where the average preferred spatial frequency for units recorded in V1 [26] was greater than in higher-order areas V2 [27] and MT [28] (Fig 2F). However, whereas the preferred spatial frequency monotonically declined for the macaque visual system, there was a U-shaped profile for the model, with the average preferred spatial frequency increasing from G_2_ to G_3_. In contrast, the mean preferred temporal frequency increased across the model’s groups similarly to the macaque visual system, with higher model groups and higher-order visual areas both showing greater selectivity to higher temporal frequency stimuli than the first model group (G_1_ vs. G_2_, t-test, *t*(87.7)=-6.07, *p*<0.0001; G_1_ vs. G_3_, t-test, *t*(31.7)=-7.32, *p*<0.0001) and macaque V1 [28], respectively (Fig 2G, 2H).

Finally, we analyzed the local connectivity of G_1_ units and found that the network replicated the results previously described for the single-layer recurrent temporal prediction model (Fig S3) [6]. Specifically, we found that short-range connections were more common for pairs of orientation-selective G_1_ units that shared a similar orientation preference (Fig S3A) and for pairs of direction-selective G_1_ units when they shared similar or opposite preferred directions of motion (Fig S3B), as is found in mouse V1 [29]. Finally, for connections between units whose receptive fields were displaced over a larger region of visual space (‘long-range’ connections), we found that the connection probability was greatest between units with a similar orientation preference whose receptive fields were aligned co-axially in visual space, again as is found in mouse V1 [30] (Figs S3C-E).

Thus, overall, the model qualitatively matched the receptive field structure, distribution of modulation ratios and the change in preferred spatial and temporal frequencies across the macaque visual hierarchy as well as the local functional connectivity rules that have been found in V1.

### Model units mirror the visual system’s hierarchy of 2D motion sensitivity

Although direction selectivity is present in V1, neurons in this area are generally unable to represent the complex motion of two-dimensional moving surfaces or patterns [31]. This sensitivity to two-dimensional motion can be probed using plaid stimuli, which consist of two overlaid full-field gratings moving in different directions (Fig 3Ai). Perceptually, the direction of motion corresponds to the average of the direction of the two components. Accordingly, representing this plaid pattern motion requires integrating the motion signals from the individual grating components such that neurons across the visual system are differentially sensitive to the two-dimensional motion of the plaid versus its individual component gratings (Fig 3Ai). In macaque V1, neurons generally respond to the motion of individual components making up the plaid pattern, whereas a proportion of MT neurons signal the overall motion of the plaid pattern [32].

**Fig 3.**
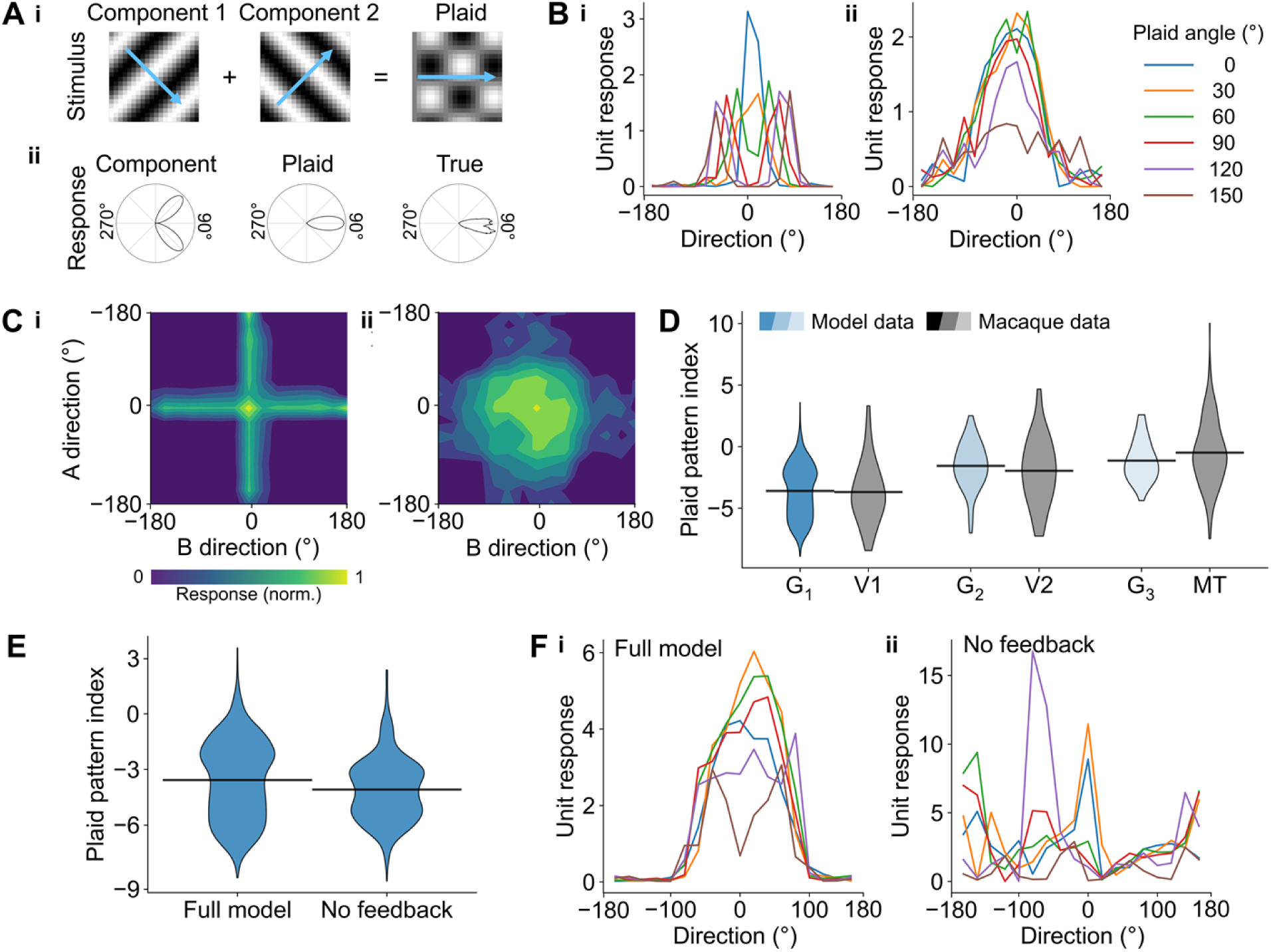
Model units mirror the visual system’s hierarchy of 2D motion sensitivity. (A) Construction of plaid stimuli through the additive combination of two grating stimuli (i). The plaid pattern index is computed by measuring the Fisher-transformed difference in the correlation between the true response of the model unit or visual neuron and an idealized component or plaid response (ii). (B) Example direction tuning curves to plaid stimuli with different plaid angles (the angular separation between the two gratings) for component- (i) and pattern-selective (ii) units. (C) Example contour plots for component (i) and pattern-selective (ii) units where the angle of each component is independently varied. (D) Plaid pattern index across the 3 model groups, G1, G2 and G3, and in macaque V1 [34], V2 [27], and MT [34]. A larger plaid pattern index indicates greater response correlation to the theoretical plaid response. (E) The mean plaid pattern index for G_1_ units is reduced when feedback in the model is abolished. (F) Example direction tuning curves for plaid stimuli across different plaid angles for the same model unit with (i) and without (ii) feedback, demonstrating a change from a pattern-like to a component-like response when feedback is abolished.

Individual model units included examples of both kinds of response to plaid stimuli. For component-selective units, the tuning curve displays two peaks corresponding to the direction of motion of the components of a plaid (Fig 3Bi), resulting in a cross-like contour plot when the direction of motion of each component is varied independently (Fig 3Ci). In contrast, pattern-selective units respond with a single peak corresponding to the overall direction of motion of the plaid stimulus (Fig 3Bii), resulting in a single response region in the contour plot (Fig 3Cii).

The plaid pattern index quantifies this preference, based on the difference in correlation between the unit’s true response and an ideal component response versus an ideal plaid pattern response (Fig 3Aii). Thus, a higher plaid pattern index indicates greater plaid selectivity. The distribution of plaid pattern index values across model groups recapitulated the hierarchy found in the macaque visual system (V1 [32], V2 [27], and MT [32]), with the plaid selectivity monotonically increasing from lower to higher groups (G_1_ mean = -3.60, G_2_ mean = -1.56, G_3_ mean = -1.12; one-way ANOVA, *F*(2, 931)=61.6, *p*<0.0001) (Fig 3D). Indeed, no significant difference was found between the means of each group and the corresponding region of the macaque visual cortex (t-test, all *p*>0.153), indicating that progressively greater selectivity for the plaid stimulus compared with the individual grating components was present across both the hierarchical recurrent temporal prediction model and the macaque visual system.

To test whether model units were specifically selective to the direction of plaid motion – as opposed to the plaid’s orientation, for example – we measured the plaid pattern direction selectivity of each pattern-selective model unit (plaid pattern index > 0). As expected, the majority (73.2%) of pattern-selective units were also pattern-direction-selective, defined as having a direction selectivity index (DSI) greater than 0.3 [6] (mean DSI=0.65). Thus, these units were specifically tuned to the direction of pattern motion.

Finally, we investigated the role of feedback in supporting pattern-like responses in model units. Although pattern direction selectivity is more weakly represented in V1, pattern-like motion responses are not entirely absent, which may result in part from feedback to V1 from higher cortical areas. Indeed, suppressing feedback from the middle suprasylvian gyrus has been shown to reduce pattern-like responses in early visual cortex of the cat [33]. In line with these experimental data, abolishing feedback in the model resulted in a significantly lower mean plaid pattern index in G_1_ (t-test, *t*(1519)=5.33, *p*<0.0001; Fig 3E), which at the level of individual model units resulted in a change from a plaid-like response (Fig 3Fi) to a component-like response (Fig 3Fii).

### Model units recapitulate feedback-dependent surround suppression

We next investigated the functional role of the model’s feedback connectivity with respect to surround suppression. Surround suppression is a hallmark nonlinearity in the responses of V1 neurons and occurs where the neural response is inhibited by stimuli extending beyond the neuron’s classical receptive field [14,15]. In the context of predictive coding, surround suppression has been interpreted as a consequence of the statistics of the animal’s natural environment [21]. As contours are generally continuous in the environment, they can be consistent with the visual system’s higher-order predictions. By contrast, short discontinuous contours violate this internal model and produce a larger response, leading to the surround suppression effect. In support of this interpretation, reducing or ablating feedback from higher visual areas to V1 reduces the magnitude of surround suppression [14,15]. Although the temporal prediction model is not trained with explicit error propagation between groups, as in predictive coding models, we asked whether ablating feedback would similarly reduce the surround suppression effect.

In line with the experimental data, surround suppression was significantly reduced when model feedback was abolished. The average G_1_ unit activity for the largest stimulus size was 24.8% smaller for the full model compared with the model where feedback was abolished (Fig 4Ai; t-test, *t*(1432)=-4.48, *p*<0.0001), which is comparable to mouse V1 data [15] (Fig 4Aii, 33.6% smaller response) though less than that for macaque V1 data [14] (Fig 4Aiii, 52% smaller response). This effect was confirmed by considering the suppression index, which describes the extent to which each unit or neuron is suppressed with increasing stimulus size. The average suppression index was significantly lower for the full model (median=51.6) compared with the model without feedback (median=19.8; Mann-Whitney U test, *U*=519075, *p*<0.0001), indicating greater surround suppression with the full model (Fig 4Bi). This effect was comparable for both mouse V1 (Fig 4Bii) and macaque V1 (Fig 4Biii), though the reduction in surround suppression was more modest in the biology.

**Fig 4.**
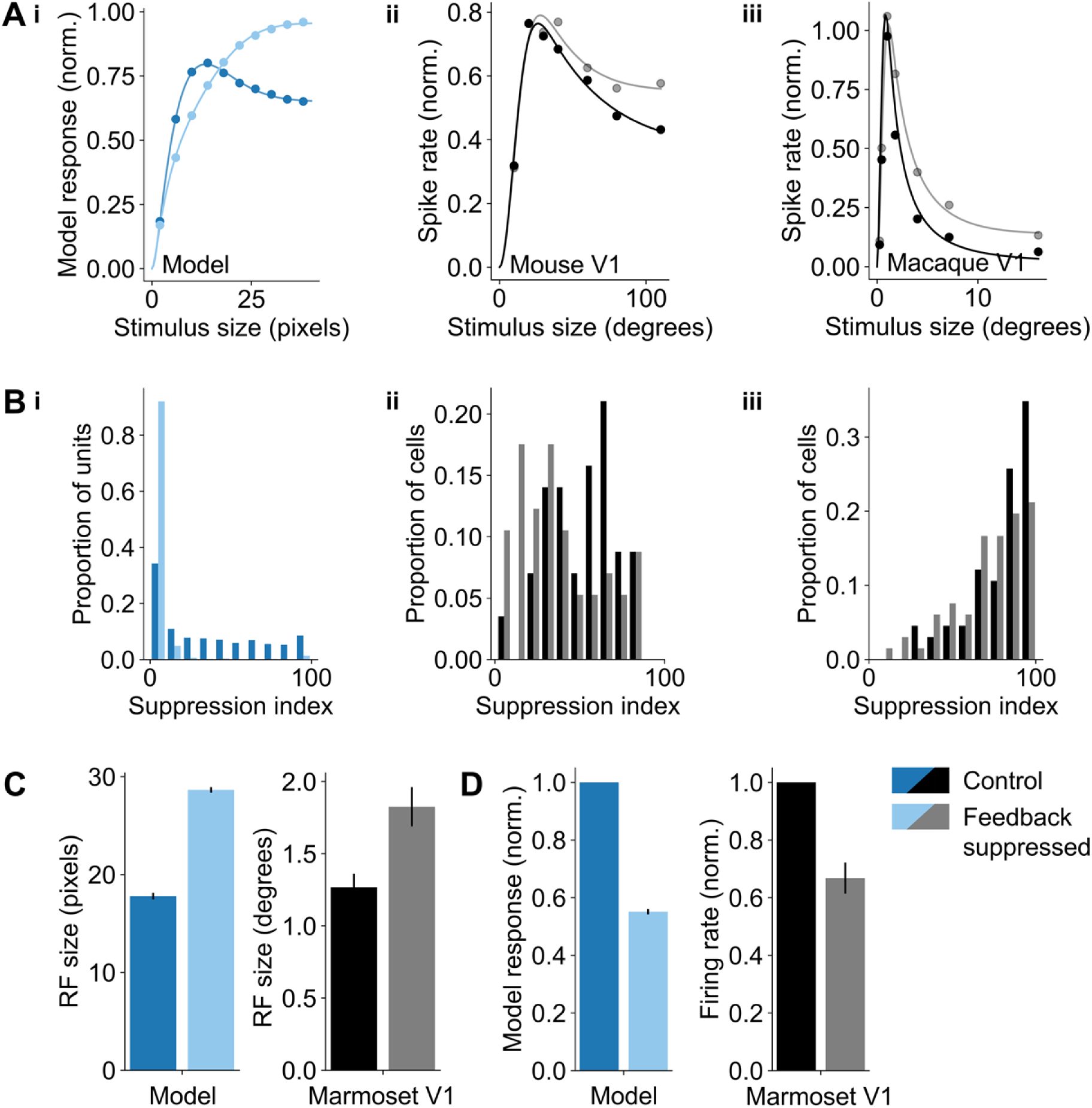
The role of feedback connectivity in surround suppression. (A) Mean response across model G_1_ units (i), mouse V1 neurons (ii) and macaque V1 neurons [14] (iii) as a function of stimulus size with and without feedback for model units or with and without inactivation of higher visual areas (mouse) or V2 (macaque). (B) Distribution of suppression index values for model G_1_ units (i), mouse V1 neurons [15] (ii) and macaque V1 neurons [14] (iii) as a function of stimulus size with and without feedback for model units or with and without inactivation of higher visual areas (mouse) or V2 (macaque). A higher index indicates greater surround suppression. (C) Receptive field size is larger when higher-order feedback is suppressed for both model G_1_ units (left) and marmoset V1 [13] (right). (D) Model responses in the classical receptive field are suppressed when higher-order feedback is suppressed for both model G_1_ units (left) and marmoset V1 [13] (right).

We further analyzed surround suppression in the model with respect to the receptive field properties with and without feedback. Specifically, we analyzed the model’s G_1_ unit responses to stimuli within each unit’s classical receptive field (defined as the maximally exciting stimulus size) as well as to stimuli in the proximal receptive field (defined as the ring of visual space area beyond the unit’s classical receptive field). In line with data from marmoset V1 [13], the average G_1_ model unit’s classical receptive field size was significantly larger when feedback was abolished (paired t-test, *t*(790)=-29.9, *p*<0.0001; Fig 4C). We next analyzed the average response of G_1_ model units for stimuli in each unit’s classical receptive field, as well as for stimuli that included both the classical and proximal receptive fields. As for marmoset V1, the normalized response was significantly smaller for stimuli within each unit’s classical receptive field when feedback was abolished (one-way t-test, *t*(790)=-48.3, *p*<0.0001; Fig 4D). However, unlike in marmoset V1, we found that model G_1_ units’ average response was suppressed for stimuli spanning the classical and proximal receptive field when feedback was abolished, whereas it increased slightly in marmoset V1 (Fig S4).

### The hierarchical recurrent temporal prediction model matches cortical stimulus representations better than other models

We next compared how well different normative models could recapitulate the response properties across the visual pathway in terms of the modulation ratio, plaid motion selectivity and the presence of surround suppression. To probe the impact of recurrency while maintaining a hierarchical representation, we compared the model to a purely feedforward hierarchical temporal prediction network. Similarly, to assess the role of the training objective, we compared the hierarchical recurrent temporal prediction model with a hierarchical recurrent autoencoder. For each model, we quantified the difference between the experimental and model distributions of each measure using the Kolmogorov-Smirnov distance, where a lower value indicates that the two distributions are more similar.

The distribution of modulation ratio values across the model groups was closer to those measured for macaque V1 and V2 neurons in the hierarchical recurrent temporal prediction than in the comparison models (Fig 5Ai, ii). The autoencoder exhibited weak bimodality among the first group of units but, unlike macaque V2 responses, was skewed towards a modulation ratio greater than one in the second group. In contrast, the feedforward temporal prediction model produced only simple-cell-like units in the first group, with complex-cell-like responses emerging only in the second group. Thus, the recurrency in the full model was required for the emergence of simple- and complex-cell-like units within the first group.

**Fig 5.**
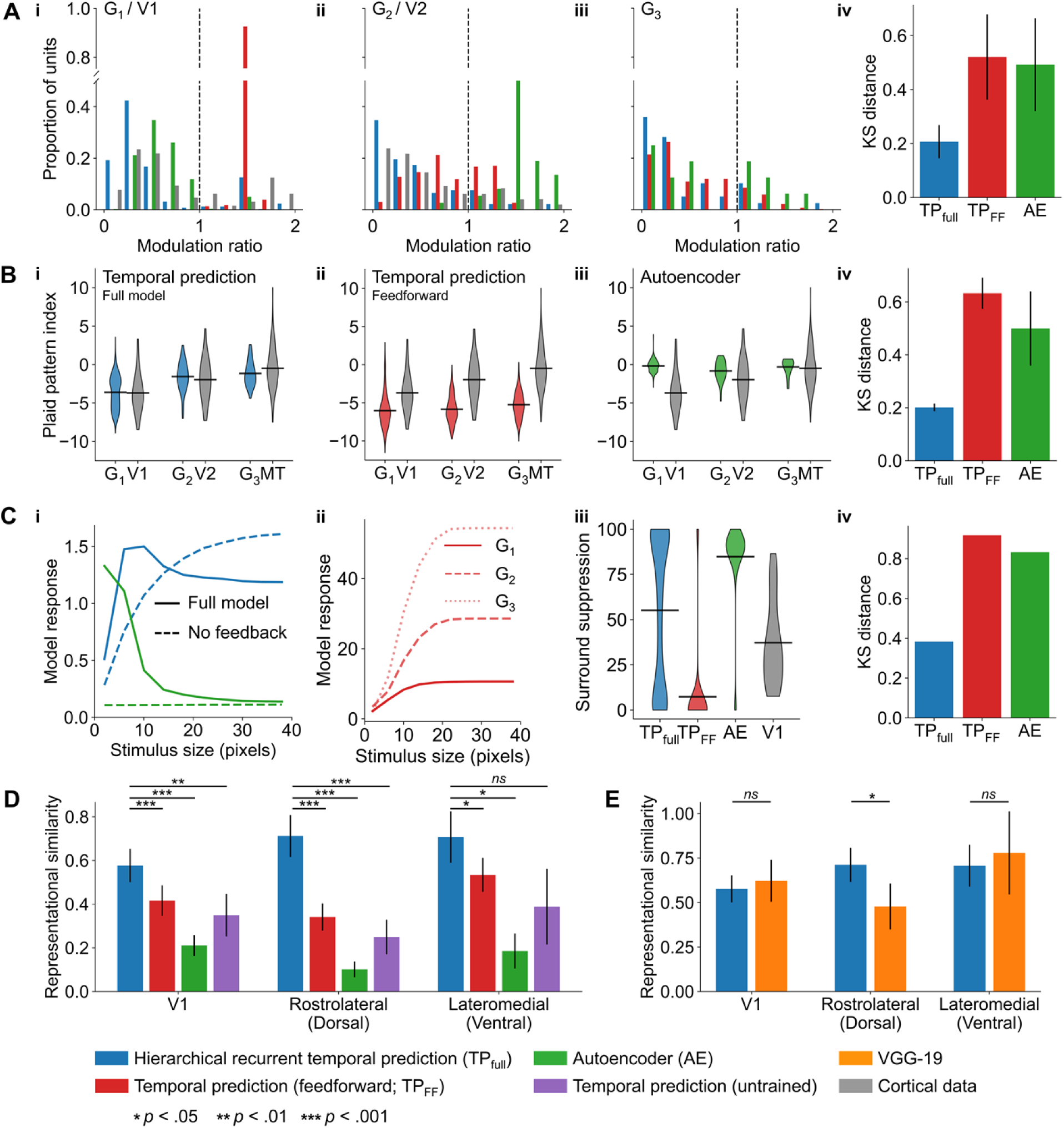
Comparison of tuning properties across the models. (A) Distribution of modulation ratio values across model units and experimental data (i: G_1_ vs macaque V1, ii: G_2_ vs macaque V2, iii: G_3_) and the Kolmogorov-Smirnov (KS) distance from the experimental data (iv). (B) Distribution of plaid pattern index values in macaque V1, V2 and MT and across groups for the recurrent temporal prediction model (i), feedforward temporal prediction model (ii), autoencoder model (iii) plus the KS distance between these models and macaque data (iv). (C) Surround suppression among G_1_ units for the hierarchical recurrent temporal prediction and autoencoder models with and without feedback ablated (i), for each group of the feedforward temporal prediction model (ii), the distribution of surround suppression index values for each model and mouse V1 (iii), and the KS distance between each model and mouse V1 (iv). (D-E) Representational similarity of the recurrent temporal prediction model relative to comparison normative models with mouse V1, rostrolateral visual cortex and lateromedial visual cortex.

A monotonic increase in the mean plaid pattern index was found across the groups in both the feedforward and hierarchical recurrent temporal prediction models, which is consistent with a hierarchical organization of global motion sensitivity (Figures 5Bi, 5Bii). However, the population mean across groups was significantly lower for the feedforward model (t-test, *t*(1790)=28.1, *p*<0.0001), which produced very few plaid-like responses. In contrast, there was relatively little variation in pattern selectivity over component selectivity as a function of model group in the autoencoder model, and the mean pattern index was overall greater than that found in macaque V1 or V2 (t-test, *p*<0.022; Fig 5Biii). Thus, the emergence of pattern-selectivity across the visual pathway was best captured by the hierarchical recurrent temporal prediction model.

For surround suppression, we first compared the mean responses as a function of stimulus size with and without feedback for the hierarchical recurrent temporal prediction and autoencoder models (Fig 5Ci). For the full models with feedback, surround suppression was greater for the autoencoder model than for the temporal prediction model, although the response for the autoencoder was very weak once feedback was abolished. In contrast, for the feedforward temporal prediction model, the mean tuning curve showed no evidence of surround suppression for any group, though the mean response increased with model depth (Fig 5Cii). These results were confirmed by comparing the distribution of surround suppression across all models (Fig 5Ciii). Although the bimodal distribution of the hierarchical recurrent temporal prediction model was qualitatively distinct from the distribution of these values in mouse V1, the mean suppression index (mean=55.1) was closer to the V1 data (mean=37.2) and the Kolmogorov-Smirnov distance was much smaller than for the other models (Fig 5Civ). The mean suppression index was larger for the autoencoder model (mean=84.8), where fully suppressed units were overrepresented, whereas only a few units showed any degree of surround suppression in the feedforward model (mean=7.3).

To further relate the networks’ learned representations to the visual system, we quantified the similarity of each model to different regions of the mouse visual cortex. For each model, we computed the representational similarity by first calculating the response similarity matrices – the correlation of model or neural activity across pairs of stimuli – before measuring the distance between these similarity matrices (Fig S5). In this way, representational similarity gives a population-level measure of model-brain alignment. For all representational similarity analyses, the reported values are for the highest performing group or layer across each network (these results did not qualitatively change when all units were grouped together for the hierarchical recurrent temporal prediction model variants) (Fig 5D, E).

In terms of brain regions, we considered mouse V1 as well as two early regions of the ventral (rostrolateral visual cortex) and dorsal (lateromedial visual cortex) visual streams. In the primate visual system, the dorsal- and ventral-streams are characterized by their relative selectivity to motion and form, respectively [35]. The mouse visual system similarly features a dorsal- and ventral-like stream [36,37] and while the distinction between these streams is less well characterized, they are differentially sensitive to motion, as in the primate visual system [38]. Accordingly, they provide an opportunity to investigate how the motion tuning of the temporal prediction model relates to the model-brain alignment in the stimulus representations across these three visual areas.

For each of these cortical regions, the representational similarity of the hierarchical recurrent temporal prediction model exceeded that of both the feedforward model (paired t-test, all *p*<0.023) and the autoencoder model (paired t-test, all *p*<0.011) (Fig 5D). We also compared the full model against an untrained variant and again found that the hierarchical recurrent temporal prediction model had higher similarity scores in all cortical areas, with significant differences between the models in V1 (paired t-test, *t*(13)=4.01, *p*=0.001) and rostrolateral visual cortex (paired t-test, *t*(12)=7.31, *p*<0.0001), but not in lateromedial visual cortex (paired t-test, *t*(9)=2.26, *p*=0.050) (Fig 5D). Given the greater motion sensitivity of the dorsal stream, we hypothesized that the hierarchical recurrent temporal prediction model would perform better as a model of a dorsal region (rostrolateral visual cortex) versus a ventral region (lateromedial visual cortex) relative to a model optimized on static images (Fig 5E). Accordingly, we compared the hierarchical recurrent temporal prediction model to a model trained for object recognition – namely, VGG-19 [39]. In support of this hypothesis, we found no difference in the representational similarity score between the temporal prediction model and VGG-19 across V1 (paired t-test, *t*(13)=2-0.70, *p*=0.499) and lateromedial visual cortex (paired t-test, *t*(12)=2.60, *p*=0.023), whereas the temporal prediction model was significantly better for the dorsal rostrolateral visual cortex (paired t-test, *t*(9)=-0.41, *p*=0.694).

Overall, these results show that the distribution of the tuning parameters examined and model-brain alignment were best captured by the hierarchical recurrent temporal prediction model.

## Discussion

The principle of temporal prediction argues that the sensory brain is optimized to represent the immediate future input based on the recent past. In line with this principle, we hypothesized that a hierarchical recurrent model optimized for temporal prediction should recapitulate the functional organization of visual cortex. With little fine tuning to the model, the network exhibited tuning properties akin to those found across several visual cortical areas, including not only the distribution of simple- and complex-like responses found in macaque V1 and V2, but also feedback-dependent surround suppression and the emergence of global motion sensitivity across those visual regions leading to area MT. Compared with alternative normative models, the response properties of visual cortical neurons were best captured by the hierarchical recurrent temporal prediction model, which similarly exhibited the closest alignment with the stimulus representations found in mouse visual cortex. Together, these results provide evidence for temporal prediction as an organizational principle across the visual cortex.

### Relation to biology

As a normative model, the hierarchical recurrent temporal prediction model represents a fairly abstract representation of the visual brain [3,40], rather than focusing on low-level mechanistic details. It is important to find a careful balance between incorporating sufficient details to make meaningful comparisons with the biology possible, while avoiding potentially redundant features that could obscure the role of the more general normative principle of interest. In the case of the current temporal prediction model, the network’s units obey Dale’s law – projecting exclusively excitatory or inhibitory connections – and incorporate both local recurrent and feedback connectivity. These fundamental properties of neural circuits are generally omitted from normative models of sensory processing, particularly those implemented as deep object recognition networks [10,41].

However, as point-like neurons, the model’s units are fairly abstract representations of true biological neurons. That is, they lack different neuronal compartments and their activity is rate based and omits spiking. Nevertheless, these neuronal properties are fully compatible with temporal prediction as a principle and implementing them in other studies has enabled the action potential firing patterns and membrane time constants of neurons to be reproduced [42,43]. Furthermore, the model is trained via backpropagation, which is generally considered to be biologically implausible [44]. However, the model itself is agnostic about the learning rule – certain elements of temporal prediction could be genetically hard-wired or arise via competitive activity-dependent mechanisms during development – and novel more biologically-plausible algorithms are being developed, which could, in principle, be applied to the current network [45,46].

### Comparison with alternative models

In the current study, we compared the hierarchical recurrent temporal prediction model to several alternative models to better understand how each model’s features related to their capacity to capture different elements of the visual system.

The feedforward temporal prediction model incorporated both a hierarchical organization and the temporal prediction objective – that is, each higher group predicted the future activity of its lower order inputs, but in contrast to the full temporal prediction model, did not include local recurrency or feedback connectivity. As shown in previous work, the distribution of model response properties in the feedforward model mirrored some aspects of the dorsal visual pathway [4]. However, plaid motion selectivity and surround suppression were considerably weaker than in the recurrent model, and the feedforward model performed worse in terms of its representational similarity to the visual cortex. Accordingly, the architectural addition of local and long-range recurrency had a large impact in improving the model’s fit to the visual system. Similarly, in terms of the model objective, the hierarchical recurrent autoencoder model performed worse on the response similarity measure and did not match the visual system’s response properties as well as the temporal prediction model. Thus, both the temporal prediction objective, as well as the model’s architecture, were important in producing a brain-like model.

Several other studies have also modeled the visual system by applying unsupervised learning principles to dynamic spatiotemporal inputs. In particular, a number of studies have investigated how well dual-stream networks can model both the dorsal and ventral streams when trained on dynamic moving visual inputs. Bakhtiari et al. [47] demonstrated that a dual-stream network trained for contrastive temporal prediction can develop dorsal stream-like representations that recapitulate some elements of motion processing, such as random dot kinematogram selectivity. Although more abstract in its form than the hierarchical recurrent temporal prediction model, their contrastive model underscores the value of temporal prediction as an unsupervised principle for modeling sensory areas. However, unlike in this study, their contrastive objective was trained to predict the *latent* state of the future frame rather than the pixel-wise future frame itself. Considering the successes of contrastive methods elsewhere [48], it will be instructive to explore this form of temporal prediction in future work. Nevertheless, their model – as is common with other contrastive networks – relies on a pretrained ResNet backbone and therefore is not truly unsupervised in the same manner as the models described in this study.

Similarly, other studies employing a dual-stream network, such as Cadieu and Olshausen [49], have described how optimizing a dual-pathway network – in their case to decompose inputs into amplitude and phase information – can produce ventral- and dorsal-like representations. After training, the optimized network produces simple- and complex-cell-like responses, as well as units with various kinds of form and motion invariance. Although their model lacks recurrency and is limited in the range of physiological properties that were investigated relative to the current study, it illustrates the utility of such dual-stream models, which could be integrated into the temporal prediction framework in future.

Finally, predictive coding represents another hierarchical normative framework, which has been very influential when applied to sensory processing. Predictive coding argues that the brain is optimized to reduce statistical redundancies by passing forward only the residual prediction errors that cannot be “explained away” by the brain’s internal model [20]. Predictive coding is inherently hierarchical in that it is organized around a series of stacked prediction modules, which – like temporal prediction – is consistent with a hierarchical view of cortical organization. The original model by Rao and Ballard [21] was able to account for non-linear effects in V1, such as surround suppression, though it was only trained on static images. In contrast, a more recent variant on predictive coding – PredNet [50] – has also been trained on dynamic natural movies. While PredNet could account for some properties such as surround suppression [51], it does not generally recapitulate low-level features of the visual cortex, such as the Gabor-like receptive fields of V1 [6], nor has it been shown to capture the hierarchical organization of the visual system.

In conclusion, we have shown that a hierarchical recurrent network optimized for temporal prediction replicates many aspects of the neuronal response properties and functional organization of different areas within the mammalian visual cortex. These findings add to the growing evidence that temporal prediction can account for information processing across the sensory hierarchy.

## Materials and Methods

### Model dataset

The model was trained using a novel dataset drawn from around 2.5 hours of wildlife videos down sampled to 25-30 frames per second. Videos were pre-processed by converting them to grayscale, resizing each clip to a width and height of 600 pixels using bilinear interpolation and applying a bandpass filter. Each video was then cropped into 1200 spatially overlapping clips of 20x40 pixels at 40 contiguous frames each. Finally, each video clip was normalized by subtracting its mean and dividing by its standard deviation. This produced a training, test and validation dataset of 1000k, 200k and 200k clips, respectively.

### The hierarchical recurrent temporal prediction model

The model is implemented as a recurrent network consisting of a series of hierarchically organized groups of units (Fig 1A). During training, each group in the model is optimized to predict the future value of its inputs – specifically, the future video frame for group one (G_1_) units or the future internal state of the lower-order inputs for each group *n* > 1. Internally, the model is implemented as a single recurrent layer, with the hierarchy defined by 1) the hierarchy of prediction whereby higher-order areas are trained to predict the future value of their lower-order inputs and 2) by the restricted internal connectivity such that each group receive inputs from and projects to its immediate higher- and lower-order groups only. This internal hidden state is then mapped to the predicted future state by a linear output layer, with the prediction error between the true and expected future state minimized during training via backpropagation. Finally, an L1 regularization penalty in the cost function ensures that only weights that contribute to network performance are retained.

More formally, the model receives as input a tensor *U* of shape *T*×*I* where *t*=1 to 40 timesteps and *i*=1 to 800 pixels as a flattened 20×40 pixel image. The network itself consists of 3 groups *g*=1 to 3, where each group *g* comprised *j*=1 to 800 units. The activity of each hidden unit *h_jt_^g^* is defined as:

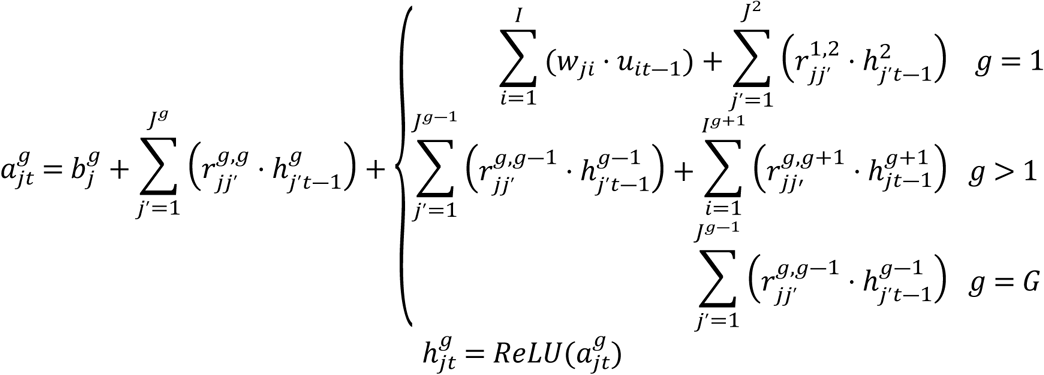

where *b_j_^g^*is the bias of group *g* unit *j*, *w_ji_* is the input weight between pixel *i* and group one unit *j*, *u_it_* _-1_ is the activity of pixel *i* at the previous timestep, *r_jj_^g,g^*is the recurrent weight between presynaptic unit *j’* in group *g* to postsynaptic unit *j* in group *g*, and *h^g^_j’t-1_*is the activity of presynaptic unit *j*’ in group *g*.

In addition, to enforce Dale’s Law whereby units have exclusively excitatory or inhibitory connections, each recurrent weight *r_jj_^g,g^* was clamped after each forward pass as:

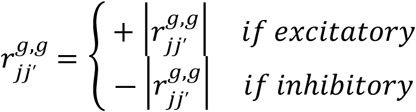

where 20% of units in each group were set as inhibitory (based on the approximate percentage of cortical inhibitory interneurons found in the brain [52,53]), with the additional constraint that inhibitory units could only project locally.

Finally, the network’s internal hidden state was mapped to the output predictions as *y_kt_^g^* for group *g*, unit *k* at time *t*, which was defined as:

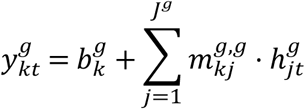

where *b_k_^g^*is the bias for unit *k* of group *g* and *m_kj_ ^g,g^*is the weight between hidden unit *h_jt_^g^* and output unit *y_kt_^g^*.

The trainable parameters in the network, *b_j_^g^*, *w_ji_*, *r_jj_^g,g^*, *b_k_^g^* and *m_kj_^g,g^*, were optimized by minimizing the loss function:

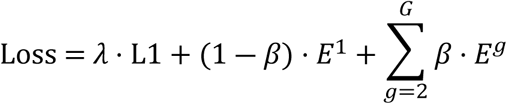

where *L1* is sum of the absolute value of all weights in the network, *λ* is a weighting parameter that describes the degree of regularization, and *β* is a weighting parameter that determines the relative contribution of the prediction error between the first and higher-order groups. Finally, *E ^g^* is the mean squared error between the true and predicted future value of the lower-order group *g-1*:

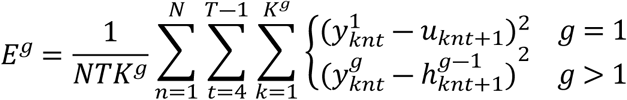

where, for group one, 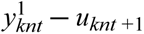 is the difference between the predicted and true future pixel value, and, for higher order groups, 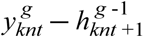 is the difference between the predicted and true future value of the lower order group. Finally, *E ^g^* is averaged across all clips *N*, time steps *T* and pixels *I*. Subscript n that indicates clip number was left off from all previous equations for brevity.

### Comparison models

Three comparison models were developed in addition to the hierarchical temporal prediction model as described above.

1. Feedforward temporal prediction model – this model was trained to maximize the same loss function as the hierarchical temporal prediction model but using a strictly feedforward architecture. The training and architectural details were adapted from previous work [4], though we only trained three stacks, and adapted the kernel sizes as detailed below to equate the receptive field sizes in the hierarchical recurrent models:

| Stack | Input size | Hidden layer size | Kernel size |
| --- | --- | --- | --- |
| 0 | 1x50x800 | 800x46x1 | 5x800 |
| 1 | 800x46x1 | 800x42x1 | 5x1 |
| 2 | 800x42x1 | 800x38x1 | 5x1 |

2. Hierarchical recurrent autoencoder – the same architecture as the hierarchical recurrent temporal prediction network but with the loss function altered to predict the activity of the current rather than future lower-order group, with an additional regularization term based on the summed activity in the network to ensure the sparsity of the network’s internal representations.

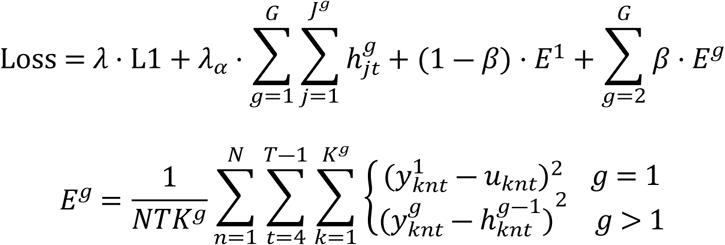

3. Untrained hierarchical recurrent temporal prediction model – the same architecture as described above, but with no training and hence randomly initialized weights.

Finally, one other pre-trained model developed by another group was also included to compare its capacity to capture the neural representation of naturalistic visual stimuli:

1. VGG-19 [39] – a deep convolutional network trained on the ImageNet database to classify images into one of 1000 categories. This network is purely feedforward and does not incorporate any form of recurrency.

### Implementation

The temporal prediction model and its variants were all implemented in PyTorch, with gradient descent performed using the ADAM optimizer set at a learning rate of α = 10^-4^. The hyperparameter λ was set as 10^-6^, as this value was close to the global minimum that minimized the mean squared error for next frame prediction on the validation dataset while producing the most biologically realistic receptive fields. The hyperparameter β was set at 0.1. Note that for Fig 1C-D, where a large number of model variants were investigated, for computational efficiency, these models were trained on a smaller 20x20 pixel dataset [6], rather than the 20x40 pixel dataset used throughout the rest of the study. All networks were trained for a maximum of 4000 epochs or until the loss function converged on a steady state.

### Data analysis

#### Receptive field estimation

Receptive fields were estimated using the response-weighted average, by taking the weighted response of each unit in the network to 200,000 frames of random Gaussian noise.

#### Unit tuning characteristics

To assess each unit’s tuning properties, we measured its response to sinusoidal gratings varying in temporal frequency, spatial frequency and orientation for 50 frames. Each unit’s preferred temporal frequency, spatial frequency and orientation was taken as the parameter or parameter combination that maximized the unit’s mean response across frames.

#### Modulation ratio

The modulation ratio was computed as F=F_1_/F_0_ where F_0_ is the mean response of the neuron to its preferred stimulus and F_1_ is the amplitude of the fitted sinusoid to the neuron’s response to its preferred stimulus.

#### Orientation and direction selectivity

Orientation selectivity was quantified via the orientation selectivity index (OSI):

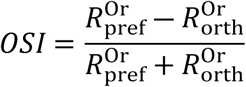

where 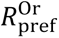 and 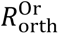 are respectively the unit responses at the preferred and orthogonal orientations for that unit. Similarly, direction selectivity was quantified via the direction selectivity index (DSI):

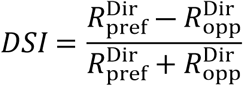

where 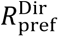 and 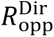 are respectively the unit responses at the preferred and opposite directions for that unit.

#### Plaid responses

Plaid stimuli were produced by summing two drifting gratings, each at half amplitude (0.5). The gratings differed by a ‘plaid angle’ – the angular separation of each plaid component (|*α* ― *β*|) around the overall direction of the plaid stimulus 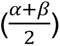. To identify the degree to which units responded to the direction of the individual plaid components versus the overall direction of the plaid stimulus, we computed the partial correlation of the unit’s response to the plaid stimulus with an idealized response assuming the unit to be entirely component or plaid selective [34]. Thus, the plaid correlation *r_p_* was defined as the correlation between the true plaid response and the predicted plaid response, controlling for the predicted component response. Similarly, the component correlation *r_c_* was taken as the correlation between the true component response and the predicted component response, controlling for the predicted plaid response. To compare across units, we computed the Z-transforms *Z_p_* and *Z_c_* of *r_p_* and *r_c_* using Fisher’s Z-transformation. This process was repeated across component separation angles of 60, 90, 120 and 150 degrees. The plaid pattern index was then taken as the difference between the average *Z_p_* and *Z_c_*values across all component separation angles. A value greater than 0 indicates greater selectivity for the plaid stimulus over its individual components, whereas a value less than 0 indicates greater selectivity to the individual components over the composite plaid stimulus.

#### Surround suppression

For each G_1_ model unit, we first fitted a Gabor function to each unit’s response-weighted average to extract the receptive field center. We then presented a drifting grating stimulus determined by the unit’s preferred orientation, spatial and temporal frequency, with a mask applied to localize the grating stimulus to the unit’s receptive field center. We recorded the response of that unit as the diameter of the mask was increased from 2 to 40 pixels. We calculate the surround suppression index based on this surround suppression tuning curve as:

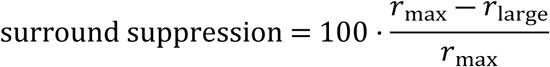

where *r_max_*is the unit’s maximum response across all grating diameters and *r_large_* is the unit’s response to the large diameter stimulus.

### Model-brain alignment

Neural data were taken from the Allen Brain Institute’s Neuropixels Visual Coding dataset of *in vivo* electrophysiological recordings of the mouse brain [54]. All recordings were pre-processed and spike-sorted by the Allen Institute. For these analyses, only data from the natural movie presentations were included (“Natural Movie One” and “Natural Movies Three”, 150 seconds total). All recordings were from wildtype mice and data from V1 (*n*=15 mice), rostrolateral visual cortex (*n*=14 mice) and lateromedial visual cortex (*n*=11 mice) were considered. In addition, only those units whose noise power to signal power ratio did not exceed 60 were included [55].

To calculate the representational similarity between each model and the neural data, we first computed the representational similarity matrix (RSM) [47,56]. For this, we binned the responses of the model or real neurons into 5-frame chunks and took the mean of each bin such that each model’s output was represented as an *M*×*N* matrix with *M* rows of stimuli and *N* columns of units. The RSM was calculated as the correlation between each pair of columns in the matrix (the response to a given stimulus *m*), resulting in an *M*×*M* RSM. The representational similarity was then calculated by taking the distance between the flattened lower triangle of the RSMs of each model and the neural data using Spearman’s correlation coefficient [57].

To compare across mice, this similarity measure was normalized by dividing by the noise ceiling for that recording, where a value ≥ 1 indicates that model similarity is as high as could be achieved once accounting for random noise [47]. The noise ceiling describes the theoretical maximum similarity that could be achieved by even a perfect model given the inherent noise present in the neural system. It was computed by randomly dividing the neural data into two halves of units and by taking the similarity of the RSMs for each half of units. This shuffling process was then repeated for 100 iterations, with the noise ceiling taken as the mean similarity value across these 100 shuffle iterations.

## Acknowledgements

This work was funded by a Wellcome Principal Research Fellowship (WT108369/Z/2015/Z) to A.J.K. S.K.-W. was supported by a studentship funded by the Nuffield Department of Clinical Neurosciences at the University of Oxford.

## Supporting Information captions

**Fig S1. Next-frame prediction performance as a function of the beta hyperparameter, Related to Fig 1**.

Next-frame prediction error increased as a function of beta for both the in-distribution (A) and out-of-distribution datasets (B).

**Fig S2. Joint distribution of modulation ratios with the orientation selectivity index (OSI) and Gabor fit *r^2^* for the first model group, Related to Fig 2**.

The simple-cell-like units (modulation ratio > 1) had higher mean orientation selectivity (A) and were better fit by the Gabor function (B) than the complex-cell-like units.

**Fig S3. Functional connectivity of group one model units, Related to** Fig 2.

(A-B) Short-range connections between model units are more prevalent for excitatory units that have similar orientation tuning (A) and direction-tuned units that have similar or opposite preferred directions of motion (B), as is also the case in V1 [28]. (C-E) In both the model and V1 [29], long-range connection probability is higher for units with similar orientation preferences when their receptive fields are located in co-axial (C) than in co-orthogonal (D) locations.

Heatmap (E) shows the normalized connection probability over visual space across differences in orientation tuning for model units.

For detailed methods, see Klavinskis-Whiting et al. [6].

**Fig S4. Model and V1 response to stimuli in the classical and proximal receptive field, Related to** Fig 4.

(A) Diagram of the classical and proximal receptive field for an exemplar neuron. (B) The full model’s average response rate is higher for stimuli spanning the classical and proximal receptive field than when feedback is suppressed. This contrasts with the data from marmoset V1, where the opposite trend is observed, such that the average neural response for stimuli in the classical and proximal receptive field is greater when feedback is suppressed [12].

**Fig S5. Schematic of the representational similarity analysis procedure, Related to** Fig 5.

## Notes

### Competing Interest Statement

The authors have declared no competing interest.

